# The Darwin–Gödel Drug Discovery Machine (DGDM): A Self-Improving AI Framework

**DOI:** 10.1101/2025.08.21.671415

**Authors:** Xuening Wu, Xinhang Zhang, Yanlan Kang, Qianya Xu, Honggang Wang, Zeping Chen

## Abstract

Despite advances such as AlphaFold and modern generative AI models, current drug discovery pipelines lack mechanisms to refine both molecules and the pipelines themselves, limiting their ability to achieve autonomous and reliable self-improvement. To address this, we present the Darwin–Gödel Drug Discovery Machine (DGDM), a self-improving artificial intelligence framework that integrates generative molecular design and evolution with adaptive meta-learning. DGDM employs a dual-loop architecture: an inner loop frames molecular optimization as a Darwinian evolutionary process guided by reinforcement learning signals, where candidate molecules generated by generative AI are evolved through search and feedback; an outer loop adaptively modifies the discovery pipeline itself. Unlike the original Gödel machine, which demands formal proofs of improvement—rarely attainable in practice—DGDM uses statistical validation to bound risk and ensure reliable progress. The framework is fully compatible with modern structural biology tools, including AlphaFold, and supports evaluation through docking, binding affinity prediction, and ADMET profiling. In a proof-of-concept study, DGDM improved the median binding affinity of candidate ligands from −4.457 to −5.422 kcal/mol while maintaining 100% drug-likeness and novelty. These results suggest that bounded-risk, self-improving AI can accelerate drug discovery by continuously refining both molecular design and discovery processes, extending the Gödel machine principle of self-improvement into biomedical research. All code is open-sourced at https://github.com/deep-geo/DGDM.

## 1 Introduction

Recent advances in AI-driven drug discovery—including diffusion-based molecular generators, graph neural networks for chemical graphs, transformer-based sequence models, and protein structure prediction systems such as AlphaFold—have greatly expanded our ability to explore chemical and protein design spaces at scale. These methods have achieved notable successes in generative chemistry, affinity prediction, and protein structure modeling. Yet most current pipelines remain architecturally static: they rely on fixed model classes, scoring functions, and search heuristics throughout the design process. Once deployed, such systems optimize only the molecules they generate, not the strategies by which those molecules are generated.

To overcome this limitation, we introduce the **Darwin–Gödel Drug Discovery Machine (DGDM)**, a self-improving AI framework that operates on two distinct levels. The *inner loop*, inspired by Darwinian evolution and enhanced by reinforcement learning, refines candidate molecules through generative modeling (e.g., large language models, diffusion architectures) and modification, with fitness assessment outcomes (e.g., docking scores, binding affinity predictions) serving as reward signals for constraint-aware selection. The *outer loop*, responsible for pipeline evolution, is inspired by the Gödel machine, which established that an AI system can self-improve indefinitely if it can verify that modifying its own code will lead to higher expected utility. In our framework, the discovery pipeline is framed as an adaptive agent that proposes modifications to models, scoring functions, or search strategies. Unlike the original Gödel machine, which requires formal proofs of improvement—often infeasible in practice—we adopt a data-driven approach where candidate updates are statistically validated. This ensures that only modifications meeting user-defined improvement criteria are accepted, with the probability of erroneous adoption bounded by a small, theoretically justified threshold. The present study demonstrates only the inner Darwinian loop, where molecules undergo modification–filter–redocking. The outer Gödel loop, with reinforcement learning and PAC-based acceptance, is not yet implemented but is outlined as a roadmap.

As a proof of concept, we implemented DGDM on a docking-based workflow, where it improved binding affinity while preserving drug-likeness and novelty. These preliminary results illustrate the potential of bounded-risk, self-improving AI to accelerate drug discovery.

### Mathematical conventions

We report results with Δ := R1 − R0 (negative Δ indicates improvement). For the acceptance analysis we use *Y* := R0 − R1 = −Δ so that larger is better. Key formulas appear in the main text; full details are in Appendix 1.

## 2 Related Work

### Protein structure prediction

Breakthroughs in protein structure prediction have transformed drug discovery. AlphaFold achieves near–atomic accuracy across diverse proteins, while newer single-sequence models such as ESMFold leverage protein language models for high-throughput prediction [1, 2]. These predictors dramatically expand structural availability, providing upstream inputs that downstream design systems—including ours—can integrate.

### Generative modeling and docking workflows

Generative AI has broadened both molecular and protein design spaces. For docking, diffusion-based models such as DiffDock frame pose prediction as a generative sampling problem, improving accuracy and enrichment [3, 4]. At the protein level, transformer-based language models (e.g., ESM-2) and structure-conditioned generators (e.g., ProteinMPNN, Chroma) enable sequence and structure design [5, 6]. In small-molecule discovery, reinforcement learning frameworks such as REINVENT [7] and graph-based approaches like the Graph Convolutional Policy Network (GCPN) [8] have been used to optimize chemical properties and synthesizability. Meanwhile, classical docking components—AutoDock Vina, Vinardo scoring, RDKit, and OpenBabel—remain widely adopted [9–12]. Our framework does not propose a new generator; instead, it integrates with existing generative pipelines, applying meta-level control to systematically enhance their performance.

### Self-improvement paradigms and cross-domain inspirations

Beyond generators themselves, recent work has explored methods that adapt and optimize the discovery process as a whole. Gödel machines formalize agents that rewrite themselves once a proof guarantees higher expected utility [13]. While such formal guarantees are infeasible in scientific pipelines, the idea motivates practical relaxations: propose edits, test rigorously, and retain only validated improvements. [14] introduced a Darwinian Gödel Machine for AI system self-evolution, illustrating how generative backbones, modification, and constraint-based filtering can drive adaptive improvement. However, direct transfer to drug discovery is impractical: unlike software, molecular design operates in noisy and continuous chemical spaces, with delayed feedback and strict requirements for safety and auditability. Our framework adapts these principles by combining constraint-driven generative search with iterative evaluation, aiming for molecules that are both chemically valid and exhibit improved docking affinity. In this sense, cross-domain inspirations from code generation and Gödel-style self-improvement provide a generalizable template for meta-evolution, which we tailor here to the molecular discovery setting.

## 3 Method

**DGDM** is organized into two nested optimization loops (Figure 1), enabling self-improvement at two levels: molecules are refined by Darwinian molecular evolution with reinforcement learning in the inner loop, while the discovery pipeline itself is adaptively reconfigured by a Gödel-style meta-optimizer in the outer loop.

**Figure 1:**
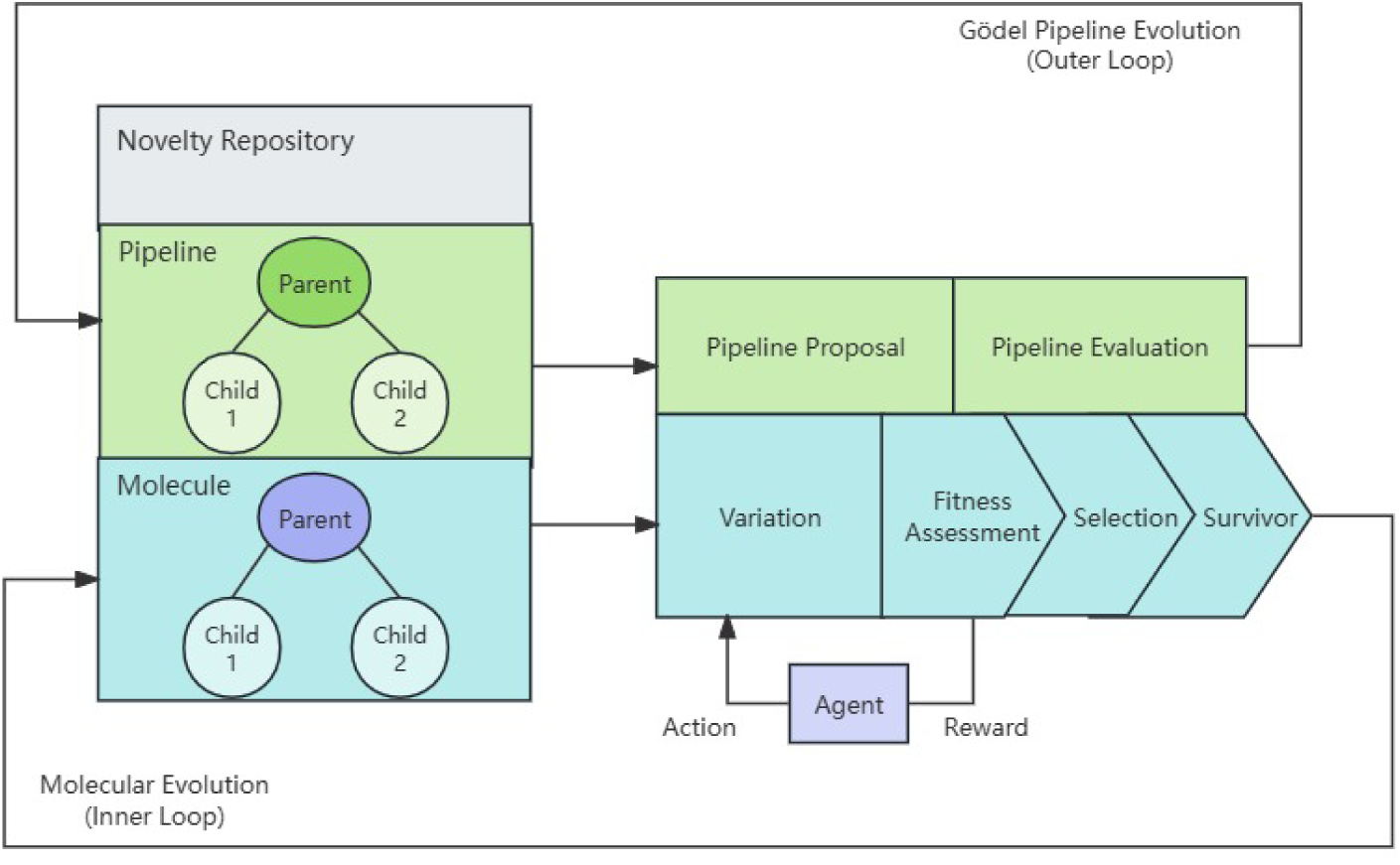
Conceptual schematic of the Darwin–Gödel Drug Discovery Machine (DGDM). The dual-loop design couples molecular evolution in the inner loop with pipeline evolution in the outer loop. Molecules are generated, modified, and optimized under reinforcement-learning–based fitness assessment, while the pipeline itself can adapt through proposed modifications to models, scoring functions, or search strategies.

### 3.1 Inner Loop: Molecular Darwinian Search

In the DGDM framework, the inner loop of molecular evolution follows a Darwinian search cycle consisting of four stages: (1) *variation/modification*, (2) *fitness assessment*, (3) *selection*, and (4) *survivor retention*. Reinforcement learning (RL) is incorporated to guide this cycle: binding affinities and constraint outcomes serve as reward signals, biasing the search toward beneficial transformations. This integration enables broad exploration of chemical space while adaptively steering evolution toward higher-quality candidates.

#### Variation / modification

Diversity in the ligand population is introduced through structural perturbations. Modifications can be realized by a range of generative AI chemistry methods, such as large language models (LLMs), diffusion models, or graph-based molecular generators. RL actions parameterize these generators—such as choosing modification operators or setting sampling conditions—so that successful strategies are more likely to recur. **Fitness Assessment:** Each modified ligand is evaluated for binding potential against the target receptor. Docking-based scoring functions, optionally coupled with structural prediction tools such as AlphaFold [1] when experimental receptor structures are unavailable, provide approximate binding free energies. These values serve as quantitative reward signals for RL.

#### Selection

Candidate molecules are ranked by predicted binding affinity, with stronger binders preferentially advanced. Selection ensures that structurally diverse yet high-scoring ligands are carried forward, maintaining evolutionary pressure toward improved binding while preserving variation within the population.

#### Survivor Retention (Constraint Filtering)

Final survivors must pass chemical validity and drug-likeness filters, including Lipinski’s rules, synthetic accessibility thresholds, and alerts for toxicity or reactive substructures. Molecules failing these constraints contribute negative feedback to RL, reducing the recurrence of unproductive modification strategies. The retained subset forms the seed population for the next generation.

Through this RL-augmented Darwinian cycle, DGDM achieves a balance between stochastic exploration and directed exploitation: exploration is realized by generative AI–driven modifications that diversify molecular structures, while exploitation is enforced through fitness-based filtering, which biases the search toward progressively higher-quality candidates across successive generations.

### 3.2 Outer Loop: Gödelian Pipeline Self-Adaptation

While the inner loop evolves molecular populations, the outer loop adapts the *pipeline* itself—that is, the configuration and ordering of operators that orchestrate molecular search. This meta-level adaptation draws on the spirit of the Gödel Machine [13], a theoretical construct for self-referential AI improvement in which a system can analyze and modify its own code to enhance performance. In the Gödel Machine’s original formulation, each self-modification required a formal proof of benefit—an approach largely infeasible in practice. We therefore adopt an empirical surrogate: candidate pipeline modifications are retained only when supported by statistical evidence of improvement, with the probability of adopting a harmful change bounded above. In this way, DGDM reflects biological evolution not only at the molecular level but also in the adaptive machinery governing its own search, combining Darwinian exploration with mathematically grounded self-improvement. The outer loop proceeds in three steps:

#### Proposal Generation

A candidate modification to the pipeline configuration is generated, for instance by using large language models (LLMs) augmented with retrieval-augmented generation (RAG) over domain-specific pipeline knowledge. Such models can suggest novel operator sequences or refinements by drawing on both general chemical reasoning and curated prior designs. As an illustration, one possible modification would be to insert a *modification–constraint refinement* step after the initial docking, where modified ligands are further substituted with fragments from a curated library and filtered by Lipinski and reactivity constraints.

#### Paired Evaluation

The modified pipeline and the baseline are applied to the same ligand set under an identical docking protocol (R1 vs. R0). A frozen evaluation harness ensures fair comparison and prevents overfitting to transient fluctuations. Significance of observed differences is first assessed using a paired *t*-test, which is standard for matched experimental designs.

#### Acceptance Test (PAC safeguard)

To guard against spurious improvements, we impose a Probably Approximately Correct (PAC)–style criterion that accepts pipeline modifications only when the risk of harm is provably bounded. Define paired improvement for replicate *i* by *Y*_i_ := R0_i_ − R1_i_ so that larger is better (*Y* = −Δ). Assume bounded scores *Y*_i_ ∈ [*a, b*] (we enforce by clipping). With the empirical mean 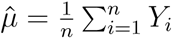, Hoeffding’s inequality yields

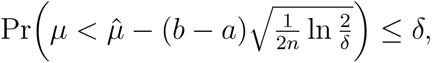

where *µ* = E[*Y*_i_]. We therefore accept a pipeline modification only if

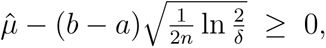

which guarantees, with probability at least 1 − *δ*, that the true mean improvement is nonnegative. In practice, a paired *t*-test is used as an exploratory diagnostic, while the PAC rule is the decisive safeguard.

### 3.3 Interaction Between Inner and Outer Loops

The *inner loop* executes a Darwinian search over candidate ligands under a fixed pipeline configuration. Its role is to generate molecular diversity, apply fitness evaluation, and filter survivors, yielding statistically meaningful performance metrics such as binding affinity and pass rate. Multiple generations of inner-loop evolution are required to obtain stable empirical evidence, and the process continues until either convergence or a bottleneck is reached. The *outer loop*, in contrast, treats the entire inner loop as a stochastic evaluator of pipeline quality. It is triggered when the inner loop meets one of two stopping conditions:

Δ_t_ *< ɛ* for *K* consecutive generations (stagnation criterion),

or

*t* ≥ *T*_max_ (iteration budget criterion),

where Δ_t_ denotes the observed improvement at generation *t*, *ɛ* is a minimum significance threshold, *K* is a patience parameter, and *T*_max_ is the maximum allowed number of inner-loop generations.

At this point, accumulated results from the inner loop are aggregated into an empirical performance estimate, which is then used to statistically test whether a proposed pipeline modification (R1) outperforms the baseline (R0). Thus, a single outer-loop decision typically relies on many inner-loop cycles until either stagnation or budget conditions are satisfied.

In this hierarchical design, the inner loop explores the chemical search space, while the outer loop adaptively refines the pipeline itself, ensuring that self-modifications occur only when supported by statistically validated evidence of improvement.

### 3.4 Evaluation Setup

Evaluation was designed to balance accuracy with reproducibility. For each ligand, we computed the median docking score across three poses to reduce sensitivity to stochastic fluctuations and outliers in the search algorithm. Both baseline (R0) and modified (R1) pipelines were evaluated under an identical docking protocol with frozen receptor preparation and scoring settings. Docking scores are interpreted here as *relative indicators* of binding propensity, not absolute free energies, in line with prior work.

Performance was assessed using four complementary metrics:

- **Binding affinity**: docking energies (kcal/mol) from Vina’s Vinardo scoring function, reflecting the predicted strength of ligand–receptor binding.
- **Score improvement**: relative change in median docking energy (Δ = R1 median − R0 median), controlling for receptor-specific biases. Negative values indicate improvement.
- **Constraint compliance (Pass rate)**: fraction of generated ligands that satisfied hydrogen-bond geometry, steric clash filters, and drug-likeness rules, ensuring synthetic plausibility.
- **Trajectory analysis**: qualitative tracing of docking outcomes across rounds to identify chemical modifications that contributed to performance gains (e.g., introduction of polar groups, repositioning of donors/acceptors).

To maximize reproducibility, we fixed all random seeds and docking parameters, and provide the full environment manifest (environment.yml) with the code repository.

### Notation and signs

Tables report Δ = R1 − R0 (negative = stronger binding under the modified pipeline). For acceptance analysis we use *Y* = R0−R1 = −Δ (larger = better) and clip *Y* ∈ [*a, b*]; in docking we set *a* = −*B*, *b* = *B* with *B* = 5 kcal/mol as a conservative range bound.

## 4 Experiments

To evaluate the feasibility of the proposed DGDM pipeline, we conducted a proof-of-concept (PoC) study on a small panel of seed ligands. The goal was to test whether generative modifications, coupled with docking-based optimization, could improve predicted binding affinities. Four chemically distinct ligands—*ASPIRIN*, *LIG3*, *LIG4*, and *PYRIDINE* —were selected to span a range of scaffold sizes and pharmacophoric features. These served as starting seeds for iterative docking and optimization. Evaluation followed the protocol in Section 3.4, with performance assessed by binding affinity, score improvement, pass rate, and trajectory analysis.

### 4.1 Docking setup

Docking simulations were performed with AutoDock Vina using the Vinardo scoring function, chosen for its balance between efficiency and sensitivity to hydrogen bonding and steric complementarity. Each ligand was docked in two sequential rounds: a baseline round (R0), where seeds were directly docked, and an optimization round (R1), where modified variants generated by the DGDM module were filtered and re-docked under identical conditions. Docking exhaustiveness was fixed at 12 across all runs, and median binding scores were reported for robustness.

### 4.2 Results

Across four seed ligands, the DGDM pipeline improved the median binding affinity from –4.457 to –5.422 kcal/mol, with all ligands passing validity filters (100% pass rate). These results demonstrate that generative modification and constraint filtering can consistently enhance predicted docking performance.

#### 4.2.1 Docking performance

Table 1 summarizes docking outcomes. All ligands improved in R1, with gains ranging from –0.807 to –1.479 kcal/mol. ASPIRIN showed the largest enhancement (Δ = −1.270 kcal/mol).

**Table 1:**
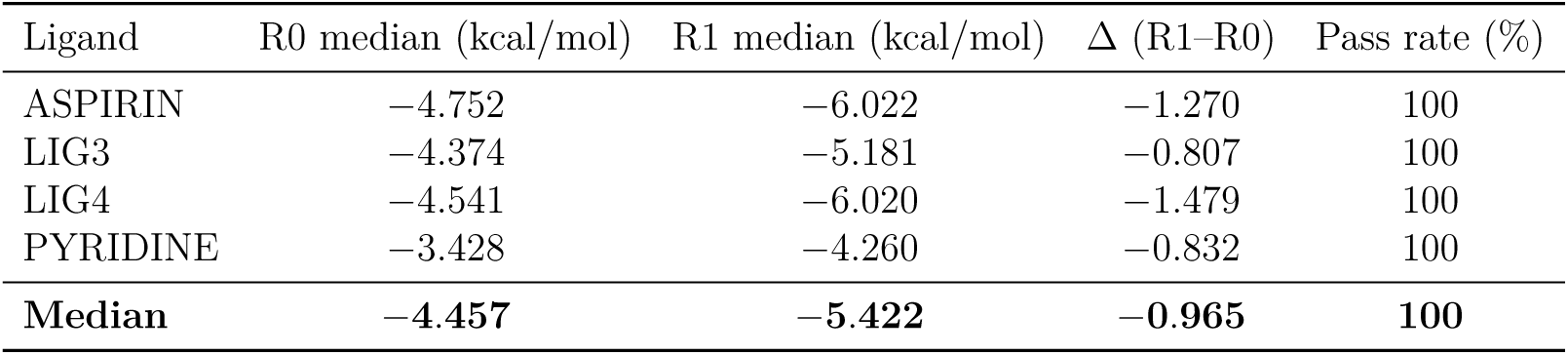
Docking evaluation results for baseline molecules (R0) and their DGDM-optimized counterparts (R1). Metrics include median docking affinity (kcal/mol), improvement (Δ = R1–R0; negative = stronger binding), and chemical validity pass rate (%).

#### 4.2.2 Representative candidate and docking trajectories

The top-performing candidate was **ASPIRIN_mut2**, which achieved a Vinardo score of –6.022 kcal/mol. In its docked conformation, the ligand aligns for hydrogen-bond formation within the binding pocket (Figure 2) while retaining the aspirin scaffold with a single substitution at the highlighted site (Figure 3). Trajectory analysis (all_rounds.csv) further indicated that overall improvements across ligands stemmed primarily from modifications introducing or repositioning hydrogen-bond donors/acceptors while reducing steric clashes, with no ligand regressing relative to its R0 baseline.

**Figure 2:**
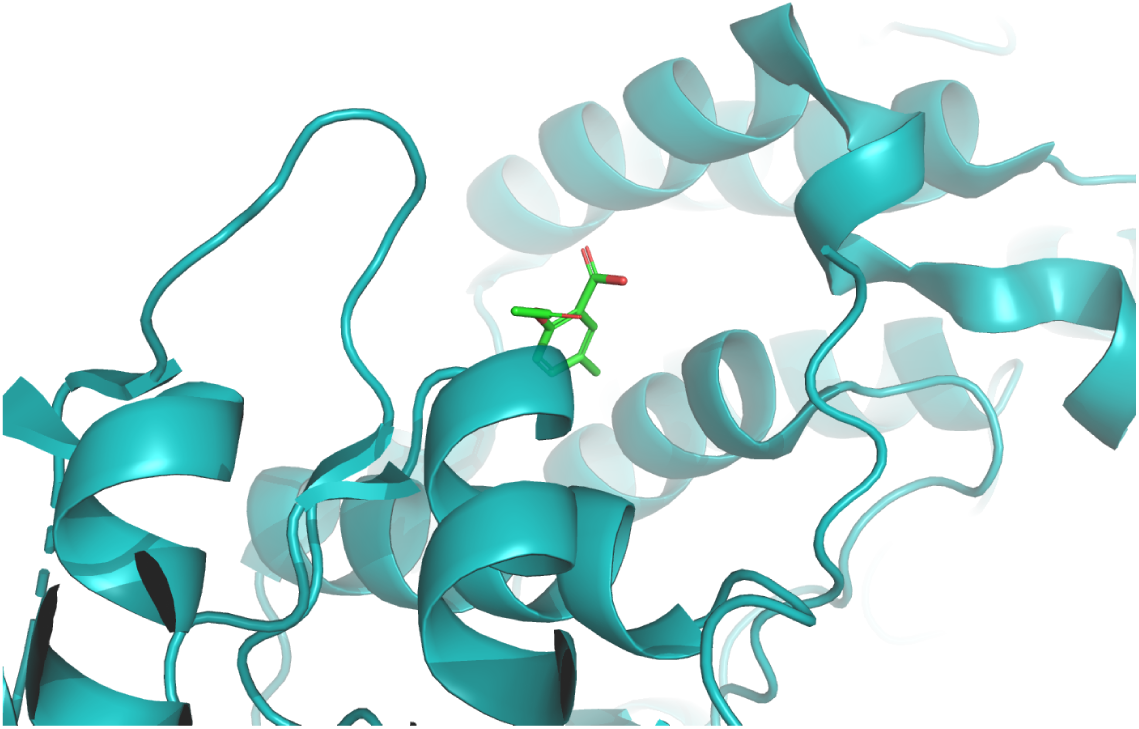
Docked pose of the DGDM-optimized candidate ASPIRIN_mut2 (green sticks) bound in the target pocket (teal ribbon). The pose illustrates improved geometric and pharmacophoric complementarity relative to the baseline seed.

**Figure 3:**
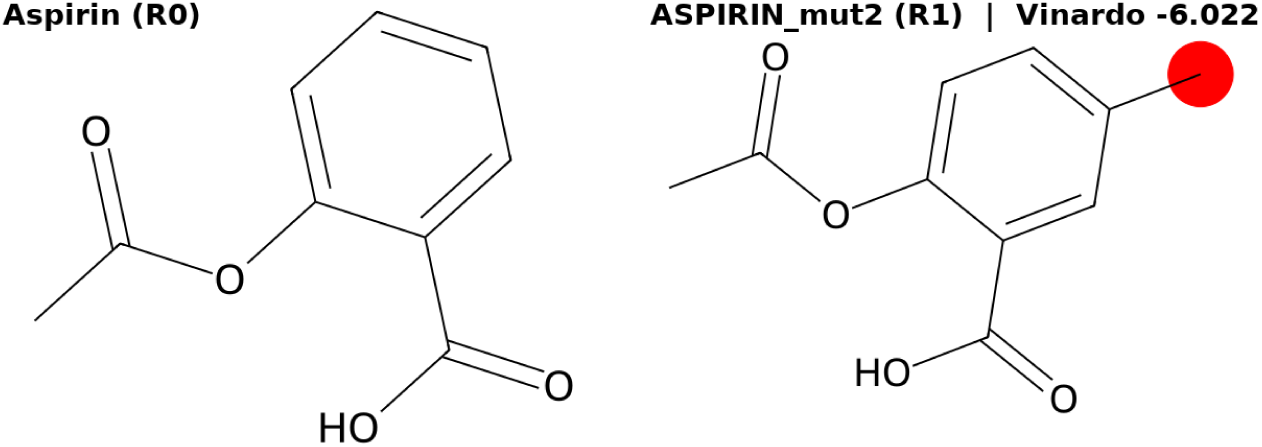
Structural comparison between baseline Aspirin (R0) and its DGDM-optimized variant ASPIRIN_mut2 (R1). The red highlight marks the modification introduced by DGDM, which enabled the improved binding pose in Fig. 2. ASPIRIN_mut2 achieved a Vinardo score of –6.022 kcal/mol versus –4.752 kcal/mol for the baseline.

## 5 Discussion

The consistent improvements observed across all four seed ligands indicate that DGDM can refine molecular structures in a systematic and reproducible manner. Unlike approaches that occasionally generate high-scoring outliers, the modification–filter–redocking cycle yielded steady affinity gains, supporting the bounded-risk design principle: structural exploration remains chemically plausible while still enabling discovery of higher-affinity variants. Even modest affinity improvements (≈1 kcal/mol) are meaningful in virtual screening, particularly when reproducible across diverse scaffolds. In this respect, DGDM compares favorably with traditional docking-only workflows, which often yield more stochastic and less reliable outcomes.

Nevertheless, the current study is a proof of concept with limitations. Evaluation was restricted to a small ligand panel and a single protein target, with docking scores serving as imperfect surrogates for binding affinity. Future work should broaden benchmarking, incorporate rescoring and molecular dynamics, and ultimately integrate experimental binding data. Importantly, this work implements only the *inner loop* of the framework. Extending to an *outer loop*—combining reinforcement learning, adaptive exploration, and wet-lab feedback—will be essential for realizing a fully closed-loop discovery engine. Experimental assays could provide the most informative rewards, grounding optimization in empirical outcomes rather than surrogate docking metrics.

Responsible deployment also requires attention. Distribution shifts may degrade performance when models encounter unfamiliar scaffolds, underscoring the need for transparent benchmarking. Reproducibility demands reporting of seeds, random states, and docking parameters. Most critically, drug discovery imposes stricter standards than many AI domains: every component—from generative operators to scoring functions—must be auditable, controllable, and governed by safeguards. Transparent reporting (e.g., model cards, audit logs) and robust governance are therefore prerequisites for safe adoption, particularly as wet-lab integration raises both scientific value and ethical stakes.

### Limitations & Scope of Preprint

This preprint reports validation of a computational framework only. The results presented here are in silico prioritizations that require downstream confirmation in wet-lab assays, such as binding experiments or cell-based screens. Experimental validation and expanded target profiling are ongoing and will be reported in the peer-reviewed version. All compounds described are computationally generated and have not been physically synthesized. Predicted biological activities are model-based and require experimental verification.

## 6 Conclusion

We presented DGDM, a Darwin–Gödel–inspired framework for molecular design that evolves both candidate structures and the optimization process itself. This proof of concept demonstrates that bounded-risk generative modification can achieve systematic affinity gains, surpassing traditional docking-only baselines. While the present study is necessarily preliminary, it highlights the feasibility of DGDM and its potential to scale into self-improving, closed-loop discovery systems that integrate computation with experimental validation, ultimately accelerating biomedical innovation with transparency and rigor.

## Appendix 1 Error-bounded acceptance in the Gödel loop

We align the analysis with the main text by defining the paired improvement

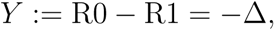

so that larger *Y* indicates better performance of the modified pipeline.

### Hoeffding bound

Let *Y*_1_*, . . ., Y*_n_ ∈ [*a, b*] be i.i.d. bounded paired improvements with true mean *µ* = E[*Y*_i_] and empirical mean 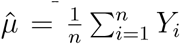. Hoeffding’s inequality gives, for any *δ* ∈ (0, 1),

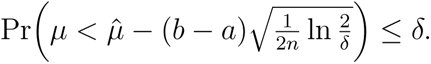

### Acceptance rule

We accept a pipeline modification when

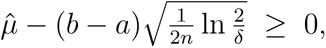

which ensures (with probability at least 1−*δ*) that the true mean improvement is nonnegative.

### Sample size requirement

To achieve an error tolerance *ɛ >* 0, it suffices to use

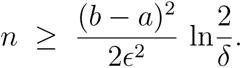

### Practical notes

i. In docking, we conservatively clip scores to *Y* ∈ [−*B, B*] with *B* = 5 kcal/mol. (ii) Variance-adaptive bounds (e.g., empirical Bernstein) can tighten radii when variability is low. (iii) A paired *t*-test can be reported as an exploratory diagnostic; the PAC rule above is the decision criterion.

## Appendix 2 Algorithmic Details

Algorithm 1 Outer Loop: Pipeline Self-Adaptation

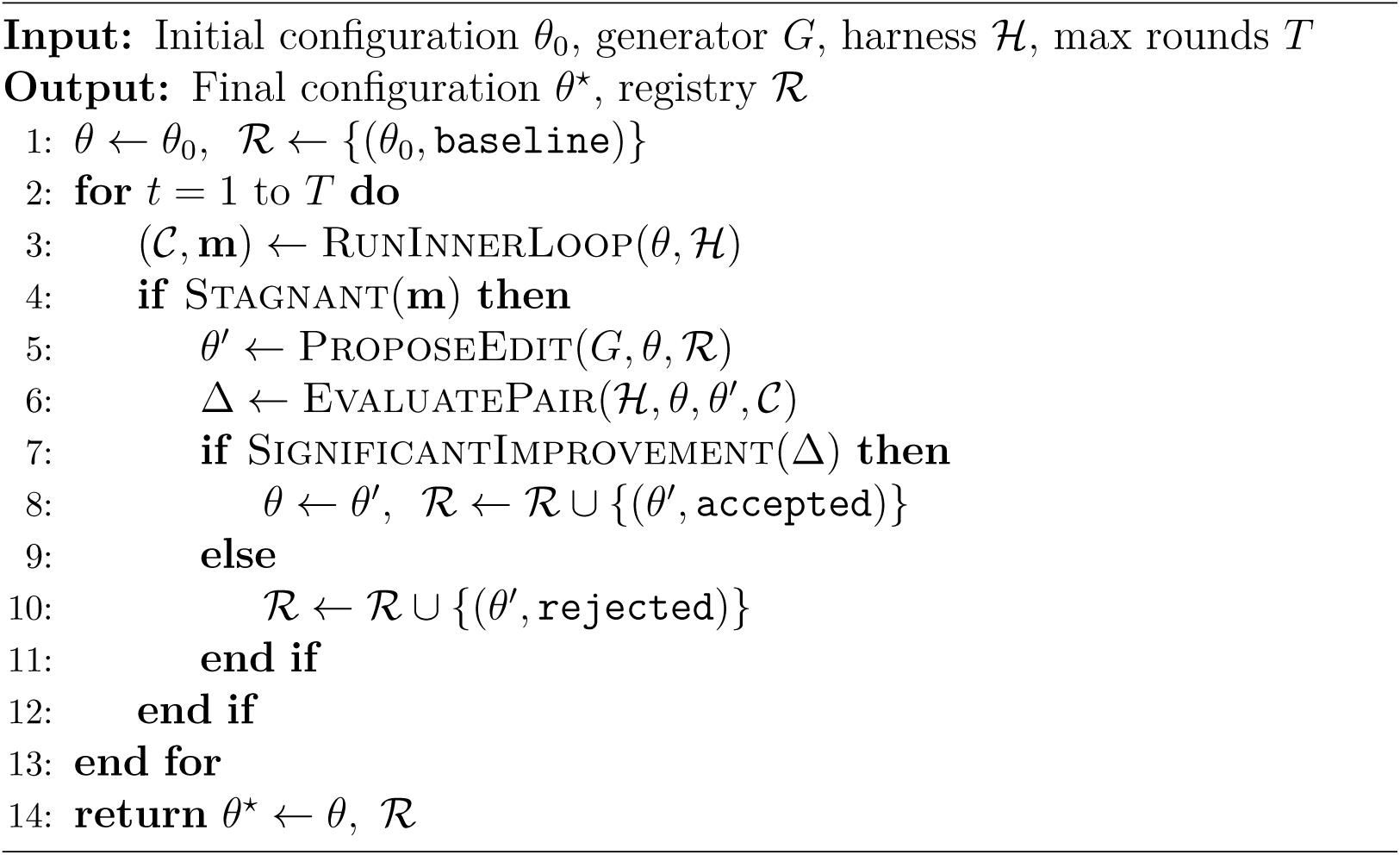

Algorithm 2 Inner Loop: Ligand Evolution (w/o RL)

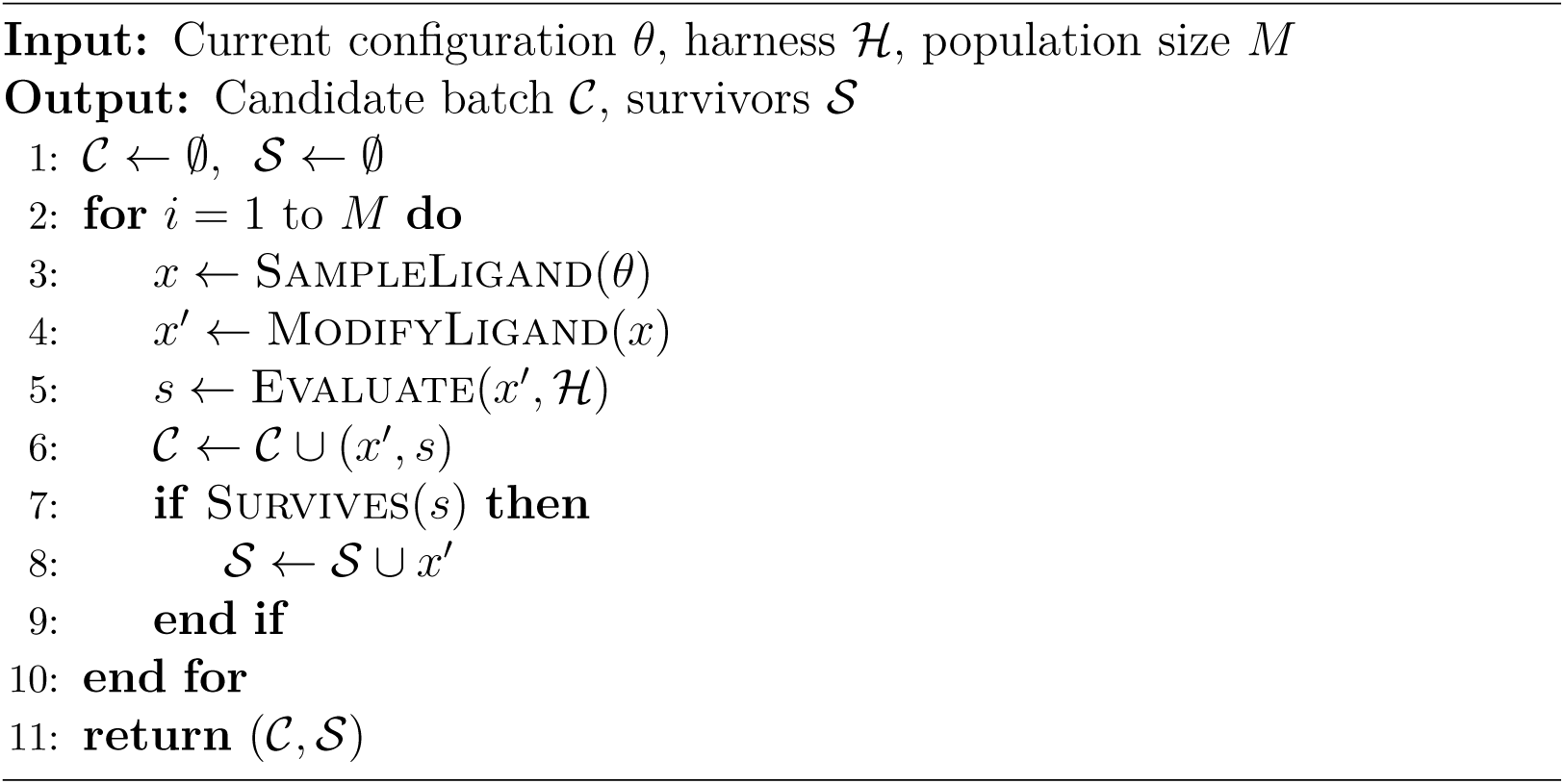

